# The TD drive - A parametric, open-source implant for multi-area electrophysiological recordings in behaving and sleeping rats

**DOI:** 10.1101/2024.03.02.582871

**Authors:** Tim Schröder, Jacqueline van der Meij, Paul van Heumen, Anumita Samanta, Lisa Genzel

## Abstract

Intricate interactions between multiple brain areas underlie most functions attributed to the brain. The process of learning, as well as formation and consolidation of memories are two examples that rely heavily on functional connectivity across the brain. In addition, investigating hemispheric similarities and/or differences goes hand in hand with these multi-area interactions. Electrophysiological studies trying to further elucidate these complex processes thus depend on recording brain activity at multiple locations simultaneously and often in a bilateral fashion. Presented here is a 3D-printable implant for rats, named TD drive, capable of symmetric, bilateral wire electrode recordings, currently in up to ten distributed brain areas simultaneously. The open-source design was created employing parametric design principles, allowing prospective users to easily adapt the drive design to their needs by simply adjusting high-level parameters, such as anterior-posterior and medio-lateral coordinates of the recording electrode locations. The implant design was validated in n = 20 Lister Hooded rats that performed different tasks. The implant was compatible with tethered sleep recordings and open field recordings (Object Exploration) as well as wireless recording in a large maze (HexMaze 9×5 m) using two different commercial recording systems and headstages.

In sum, presented here is the adaptable design and assembly of a new electrophysiological implant facilitating fast preparation and implantation.

## INTRODUCTION

The multi-area nature of brain interactions during wake and sleep makes it difficult to exhaustively study the ongoing physiological processes. While approaches such as functional MRI (fMRI) and functional ultrasound (fUS) allow to sample brain activity from whole brains^1, 2^, they exploit neurovascular coupling to infer brain activity from hemodynamic activity, limiting their temporal resolution^2^. In addition, fMRI requires placement of the research subject in an MRI scanner, prohibiting experiments with freely moving animals. Optical imaging of calcium dynamics with single or multiphoton imaging enables celltype-specific recordings of hundreds of neurons simultaneously^3^. However, head-mounted microscopes such as the Miniscope^3^, which do allow freely moving behavior, are usually limited to imaging superficial cortical areas in intact brains^4^. While the diameter of their field of view on the cortex can be in the order of 1mm, the space requirements of these head-mounted microscopes can make it difficult to target several, especially adjacent, areas. Therefore, to capture multi-area brain dynamics in wake and sleep accurately, extracellular electrophysiology, recorded with electrodes implanted in the brain areas of interest, is one of methods of choice due to its high temporal resolution and spatial precision^5^. In addition, it allows characterization of sleep dynamics in animals compatible to analyses obtained from human EEG, increasing the translational value of this method^6^.

Classically, studies recording brain activity with extracellular electrodes have employed individual wire electrodes or electrode bundles, such as tetrodes^7^. State-of-the-art probes such as the Neuropixels probe^8^ allow to target several areas simultaneously, given that they are aligned on an axis that does allow implanting the probe along that axis without impairing the animal. However, accurate simultaneous recordings of multiple, spatially separated areas still remain challenging, with existing methods being either costly or time intensive.

In recent years, additive manufacturing methods such as stereolithography have become broadly available. This allowed researchers to develop novel electrode implants adaptable to their experimental requirements^9^, for example simplified repeatable targeting of multiple brain areas. Frequently, these implant designs are also shared with the academic community as open source hardware, allowing other researchers to adapt them to their own purpose. The degree of adaptability of specific implants varies both as a result of how the implant is designed and how it is shared. Parametric modeling^10^ is a popular approach in computer-aided design, in which different components of the design are linked by interdependent parameters and a defined design history. Implementing a parametric approach for designing implants increases their reusability and adaptability^10^, as changing individual parameters automatically updates the complete designs without the need for complex re-modeling of the design. A consequential necessity is that the design itself is shared in an editable format that preserves the parametric relationships and design history. File formats that only represent geometric primitives, such as STL or STEP, make subsequent parametric modifications of published models unfeasible.

While tetrode hyperdrives^11–13^ enable recordings from dozens of tetrodes, their assembly and implantation are time intensive as well as quality being largely dependent on the skill and experience of the individual researcher. In addition, they usually combine the guide tubes that direct the recording electrodes to their target location, in one or two larger bundles, therefore limiting the number and spread of areas that can be targeted efficiently.

Other implants^14, 15^ expose the complete skull and allow for the free placement of multiple individual microdrives that carry the recording electrodes. While placement of independent microdrives^16^ during surgery time maximizes flexibility, it increases surgery time and can make it difficult to target multiple adjacent areas due to the space requirements of the individual microdrives. In addition, while the implants are open source, they are only published as STL files, making modification difficult.

An example of a drive with a more inherent parametric philosophy is the RatHat^17^. By providing a surgical stencil that covers the whole dorsal surface of the skull, it allows precise targeting of multiple brain targets without the use of a stereotactic frame during surgery. Multiple implant variations for cannulas, optrodes or tetrodes are available. However, while the drive is free to use for academic purposes, it is not published open source, creating a hurdle for researchers to evaluate and use the implant.

Presented in this article is the TD drive (see Figure 1), a novel 3D-printable implant for extracellular electrode recordings in rats. The TD drive aims to overcome some of the drawbacks of existing solutions: it allows to target multiple brain areas, mirrored across both hemispheres, with independent wire electrodes simultaneously. Due to its simple design, it can be assembled in a few hours at a relatively low costby less experienced researchers. The TD drive is published open source, in easily modifiable file formats to allow researchers to adjust it to their specific needs. Incorporating a parametric 3D modeling approach from the beginning of the TD drive’s design process allows the parameters necessary to be changed to be abstracted: to change target locations, researchers can simply edit the parameters representing their dorsoventral and anteroposterior coordinates, without the need for re-designing the drive themselves. The files to modify and manufacture the TD drive can be found at https://github.com/3Dneuro/TD_Drive.

**Figure 1:**
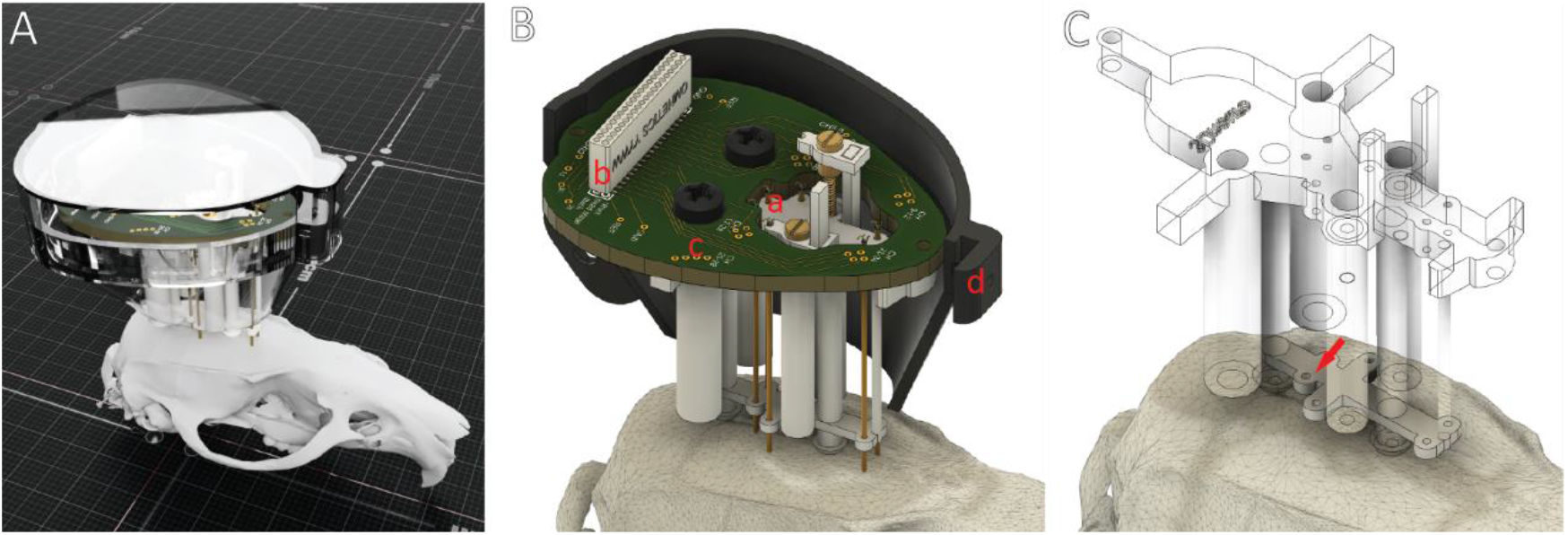
Overview of the TD drive. **A**. Rendering of a TD drive with protective cap. **B**. Rendering with inner parts shown. The TD drive features (**a)** multiple, parametrically adjustable recording locations for fixed and moveable electrode wires, an EIB with **(b)** a high-density Omnetics connector compatible to common tethered and wireless data acquisition systems and **(c)** an intuitive channel mapping optimized for recordings with Intan/Open Ephys systems and **(d)** a cap to protect the implant during tethered recordings and when no headstage is connected. **C**. A guide stencil on the bottom of the TD drive facilitates placement of guide cannulas and serves as a redundant verification of implant locations during surgery.

The implant design was piloted in n=4, validated in n = 8 and confirmed in n=8 Lister Hooded rats that performed different tasks. The first 4 animals were used to develop the drive and adjust parameters. Then a full pilot was run with 8 animals (shown in results). A second cohort of 8 animals was run and included in the implant survival analysis. The implant was compatible with tethered sleep recordings and open field recordings (Object Exploration) as well as wireless recording in a large maze (HexMaze 9×5 m) using two different commercial recording systems and headstages. The two cohorts of 8 were recorded with two different acquisition systems – tethered for longer sleep recordings and wire-less for large maze exploration recordings. We can conclude that this simple wire-drive allows for long-running experiments with larger cohorts by less experienced researchers to enable sleep stage analysis as well as oscillation analysis in multiple brain areas. This is in contrast to most electrophysiology implants to date that due to difficulty and time-intensity allow for smaller animal cohorts and usually need very experienced experimenters. However, with this drive no individual neuron activity can be recorded, thus the use is limited to investigations of LFP and summation activity.

## PROTOCOL

The present study was approved by the Central Commissie Dierproeven (CCD) of the Radboud University and conducted according to the Experiments on Animals Act (protocol codes: 2020-0020-006 & 2020-0020-010). Male Lister Hooded rats (Charles Rivers Laboratories) of 9-12 weeks on arrival were used.

### 1. Adjust and create 3D models and EIB data

1.1. Open the drive body design in Autodesk Fusion 360. Click on “Change Parameters” under the “Modify” tab. Adjust the coordinates for the first recording location by entering the anteroposterior coordinate in anteroPosteriorSite1 and the mediolateral coordinate in medioLateralSite1. You can adjust the diameter of the hole for the guide tube or electrodes by adjusting diameterSite1. Repeat for recording location 2 and 3. The model design will be automatically adjusted. NOTE: The three locations used for the current protocol are hippocampus (HPC), containing moveable wire bundles, and prefrontal (PFC) and retrosplenial (RSC) cortex, both with fixed wire bundles.
1.2. Export the updated drive body by right clicking on it in the browser and selecting “Save As Mesh”. Select type “STL (binary)”, units “mm”, and refinement “high”.
1.3. The size of the required protective cap and EIB might change depending on the parameters you chose for target locations. Select either the prepared STL files for the regular cap or, if needed (for example when targets are very lateral), the prepared STL files for the large caps.
1.4. Depending on which cap you chose, select either the regular or large EIB for production. The Gerber production files for both EIBs are provided as zip archives that can directly be sent to a manufacturing service.

### 2. Print the 3D models & Manufacture the EIB

NOTE: A Formlabs Form 3 SLA printer was used for producing the parts. When using different printers or outsourcing the production, different, comparable resins for producing the parts might need to be used.

2.1. Print the drive body and shuttles using stereolithography with a high resolution in a regular or biocompatible resin (e.g., Clear, Black or White resin) with 25 μm layer height. Print the parts for the cap with a strong and sturdy resin (e.g., Tough 2000).
2.2. Either manufacture the EIB in-house or have it produced by an external service provider such as JLCPCB. Solder the high-density connector (see Table of Materials) to the EIB using SMD soldering techniques. When not experienced in soldering fine electronic components, it is recommended to have the soldering done externally, for example at the university’s electronics workshop or a commercial supplier. Reinforce the soldered high-density connector by applying 5-minute epoxy (see Table of Materials) around the connector. Be careful not to cover the holes for the electrodes with epoxy.

### 3. Post-process 3D printed body

NOTE: Cap and shuttles should not need post-processing. Depending on the quality of the 3D prints, they might need to be lightly sanded or have leftover support traces removed. When sanding and drilling, take care not to break the walls of the drive body. If necessary, clean post-processed parts with isopropanol and a soft cloth, and/or compressed air.

3.1. Drill out the holes for the guide tubes on top and on the bottom of the drive body with a 0.5 mm drill bit (see Table of Materials) mounted in a pin vise (see Table of Materials). This ensures that the dimensions are correct and consistent across sites.
3.2. Drill out the two countersink holes (a in Figure 2E) on the drive body for the shuttle brass insert using a 2mm drill bit (see Table of Materials) in a pin vise.
3.3. Clean the countersink holes from drilling debris with compressed air. Then tap the guide holes for the shuttle screws, which are the extension of the countersink holes, with an M1 tap (see Table of Materials). Perform the tapping in two or more iterations, cleaning off debris from the tap and hole in between iterations. Optionally lubricate the tap with a drop of mineral oil (see Table of Materials).
3.4. Clean the drive body from drilling and tapping debris with compressed air.

### 4. Shuttle assemblies

4.1. Slide a 3D printed shuttle onto an M1×16 screw (see Table of Materials). Use a M1 brass insert (see Table of Materials) to hold the 3D printed shuttle in place. The shuttle needs to be able to freely rotate without moving up or down after the insert is placed. CAUTION: The following steps contain burn hazards (soldering). Depending on the solder and soldering flux used, they might include exposure to respiratory irritants and lead. Always wear eye protection when soldering (as solder can splutter) and follow appropriate guidelines for safe handing of potential harmful substances, including appropriate ventilation of the workspace to extract soldering fumes. Follow your local regulations and operating procedures or consult available material online^18, 19^.
4.2. Using a small amount of soldering paste (see Table of Materials), solder the brass insert to the screw. Take care not to overheat the insert and screw in order to not melt the 3D printed shuttle. Depending on the resin used for 3D printing the shuttle, a small amount of melting (and subsequently, the shuttle sticking to the insert) is hard to avoid. NOTE: When using stainless steel screws, solder flux (see Table of Materials) might be required. It is recommended to use brass or machine steel screws, as those are easier soldered.
4.3. After the shuttle assembly cooled down, gently rotate the 3D printed shuttle multiple times around the screw. If the shuttle fused to the insert during soldering, this should release it. Ensure that the shuttle can rotate freely and that it does not wobble. If it does, discard the shuttle assembly and start a new one. Carefully try to rotate the brass insert. If it rotates with respect to the screw, repeat the soldering process.

### 5. Assemble the drive

5.1. Cut polyimide tubes (see Table of Materials) to approximately 25 mm length, but at least long enough to extend through the complete drive body.
5.2. Insert the polyimide guide tubes into the drive body. Each tube should be inserted through one hole on the top of the drive and the corresponding hole in the guide stencil on the bottom of the drive (d in Figure 2E). The tubes should be inserted until they are flush with the top of the drive body.
5.3. Using a thin needle or toothpick, apply a small amount of liquid cyanoacrylate glue (see Table of Materials) to the holes at the top of the drive body to fix the guide tubes in place. Apply the glue from the bottom side of the body to avoid glue running into the guide tubes. The glue will be drawn into the space between drive body and guide tube by capillary forces and thus connect the two.
5.4. Apply a small amount of cyanoacrylate glue to the interface between guide tubes and the guide stencil at the bottom of the drive body. Again, take care not to clog the guide tubes with glue. Let the glue dry for a few minutes. The exact amount of time necessary depends on the drive material and clearance between drive body and guide tubes. Generally, 5-10 minutes should be sufficient.
5.5. Turn the drive body upside down and cut the polyimide guide tubes on the bottom so that they extend approximately 1 mm beyond the center pedestals of the drive body (c in Figure 2E and figure S2c). In this configuration, the end of the guide tubes will be flush with the brain surface at implantation. NOTE: the drive was developed to target deeper areas of the brain. If superficial cortical areas are targeted, shorter polyimide guide tubes might be necessary to not injure the brain surface in case of initial brain swelling.
5.6. Insert two shuttle assemblies in the drive body. While screwing them into the tapped guide holes, make sure that the screws are parallel to the shuttle guides (b in Figure 2E) extending the drive body. Use your fingers to gently align the shuttles with the shuttle guides.
5.7. Fully screw the shuttles down into the countersink holes to verify that the brass inserts of the shuttle assembly do not get stuck in the drive body or collide with the polyimide guide tubes. Do not over-tighten the shuttle in the drive body – this can destroy the trapped threads on the drive body and the solder connection of the shuttle assembly. If a shuttle assembly gets stuck, remove it completely and check whether the solder connection loosened. In that case, use a new shuttle assembly. If the shuttle assembly collides with a guide tube, shorten the guide tube so that it does not extend beyond the drive body.
5.8. Screw the EIB to the drive body with M2.5×5 polyimide screws (see Table of Materials). Apply a few drops of cyanoacrylate glue between the drive body and EIB. Make sure to not clog the through holes for electrode connection.

### 6. Prepare the protective cover

6.1. Insert a stainless steel M2 nut (see Table of Materials) into the extrusion on the left cap half and fix it with cyanoacrylate glue.
6.2. If necessary, drill out the hole at the front of the left cap with a M1 drill bit (see Table of Materials) in a pin vise. Tap the hole on the front of the right cap half with an M1 tap.

### 7. Prepare wire electrodes

7.1. Prepare two metal plates (see Table of Materials) as surface for creating the electrode wire bundles. The plates serve as a flat, stable, but moveable surface on which the wire bundle assembly, gluing and cutting will take place. Attach plotting paper (see Table of Materials) to the first plate and not too sticky painters’ tape (see Table of Materials), with the sticky surface pointing up, onto the second plate.
7.2. Three out of the four wires in the HPC bundles will be cut at a 60-degree angle to create an offset in the dorsoventral direction. This will allow placing a wire above, in, and below the hippocampal pyramidal layer, respectively. To facilitate the cut, draw a clear line with an angle of 60 degrees on the plotting paper (60-degree line).
7.3. For each HPC electrode bundle, cut 4 pieces of electrode wire with a length of 4.5 cm each. For each PFC and RSC electrode bundle, cut 4 pieces of electrode wire with a length of 3.5 cm each.
7.4. Gently pick up 4 wires by touching them with a fingertip (they will stick to it) and place them as close as possible next to each other on the painters’ tape. Make sure to not put them on top of each other.
7.5. Under a microscope, use a forceps to place the wires together as close as possible. Apply a thin layer of liquid cyanoacrylate glue to the first 2 cm of the top of the bundle. For the HPC bundle, glue > 2 cm and < 3.5 cm of the wire. Wait for the glue to dry.
7.6. Touch the wires gently with some forceps under a microscope. If they do not separate, they are glued correctly. As a sanity check, you should see the layer of glue shine under the illumination of the microscope.
7.7. Once fully dried, remove the wire bundle from the tape and transfer it to the plate with the plotting paper. Under a microscope, check the wire bundle for excess glue on top or on the sides, and remove it carefully with a scalpel blade.
7.8. For the RSC bundles, make a straight cut at the bottom of the array, perpendicular to the direction of the wires.
7.9. For the HPC bundles, place the array on the plotting paper so that it intersects the 60-degree line, and use the line as a guide to make a cut angled 60 degrees to the direction of the wires. Then, use a scalpel blade to carefully split the shortest of the 4 wires off the bundle. Cut the wire perpendicular to the wire direction, shortening it approximately 0.75 mm compared to the second-longest wire in the bundle.
7.10. For the PFC bundles, split the bottom of the array in two 2-wire bundles. Make sure that the two wires each are glued together well. Shorten one of the 2-wire bundles by 1 mm by cutting it perpendicular to the wire direction. See Supplementary S1 (bottom) and S2(a) for images of the cut wire bundles.

### 8. Prepare ground wire and EEG wires

8.1. Push at least 10 of the SIP/DIP pins out of a 1.27 mm pitch interconnected SIP/DIP socket strip (see Table of Materials).
8.2. Cut 2 pieces of 6 cm length for the ground (GND) wire (see Table of Materials). Cut 8 pieces of 6 cm length for the EEG wire (see Table of Materials). Use a scalpel blade to carefully remove some of the insulation from both ends of all wires.
8.3. Place a M1×3 stainless steel screw (see Table of Materials) in a third hand, leaving as much space as possible accessible below the screw head. Wrap a de-insulated side of a GND or EEG wire around the shank of the screw, just underneath the head of the screw.
8.4. Apply a small amount of solder flux with a small needle or toothpick. Solder the wire to the screw. Make sure to not accidentally clog the slot of the screw head.
8.5. Place a SIP/DIP pin in the third hand so that the female side is accessible. Insert the de-insulated part of the opposite side of the wire into the SIP/DIP pin. Apply a small amount of solder flux and solder the wire to the pin.
8.6. Remove the soldered screw-wire-assembly from the holder. This assembly will be implanted on the skull during implantation surgery.
8.7. Place another SIP/DIP pin in the holder, rotated 180 degrees (i.e., male side accessible). Apply a small amount of solder flux and solder a de-insulated side of the other wire to the male side of the pin.
8.8. Remove the soldered wire-pin assembly from the holder. This assembly will be later connected to the EIB, and the screw-wire assembly as well as wire-pin assembly will be connected to each other during implantation surgery using their two pins.
8.9. To reinforce the soldered connections, apply a small amount of cyanoacrylate glue to the connection between wires and pins.
8.10. After the glue has dried, verify that the SIP/DIP pins of the two assemblies can be connected smoothly. Use the check continuity option of a multimeter to verify that there is a continuous connection between the screw and the de-insulated wire end of the wire-pin assembly when both assemblies are connected. Optionally, color code each set of wires with nail polish (see Table of Materials) to simplify correct connection during implant surgery.

### 9. Load the wire bundles into the drive

9.1. Attach the drive to a holder. Be careful not to apply too much pressure to the EIB or damage the high-density connector in this step.
9.2. Once the drive body is in a stable position, take one of the wire bundles and carefully slide it into the respective polyamide tube, either using by hand or a pair of fine forceps. Make sure the wire array is placed in the correct orientations (e.g., for the PFC array the two longer wires of the array should be facing medial) and be careful not to bend the wire array.
9.3. Repeat the last step for all other wire bundles.
9.4. Use a thin forceps to grab one of the wires and carefully bend it towards the hole you want to insert it in. Once inserted, use a gold pin (see Table of Materials) to pin it into the EIB hole. Repeat this for all wires of the bundle and for all of the bundles. Make sure that, during this stage, the wires make a nice loop above the EIB (this way there is still room to move the bundle up and down the polyamide tube in order to adjust the length at the bottom of the tube) and that the array that is sticking out from the bottom of the polyamide tube does not become accidently bent. Make sure to note down which wire of each wire bundle connects to each channel on the EIB. See Supplementary Figure S1 for an elaboration on the channel mapping of the TD drive. NOTE: Alternatively, after loading of each wire bundle (step 9.2), you can directly connect the wires to the EIB (step 9.4) and then proceed with step 9.2 + 9.4 for the remaining wire bundles. This can be varied based on the experimenters’ personal preferences. See Figure S2b for an example of a loaded TD drive.
9.5. Adjust the length of the wire bundles to correctly target the recording locations by gently pushing or pulling the wire bundles in or out of the guide tube (see Figure S2d). As the guide tubes are cut to be flush with the brain surface, the distance to which a wire bundle extends beyond the guide tube corresponds to the dorsoventral location of the target area. The moveable HPC wire bundles should be flush with the bottom of the guide tube, the fixed RSC bundles should extend 1.5 mm and the fixed PFC bundles should extend 3.5 mm beyond the guide tubes. When pushing or pulling the wire assemblies, be careful not to pull individual wires out of the EIB on the top or to bend the bottom of the wire bundle.
9.6. When the fixed wire arrays (RSC and PFC) are aligned, apply a small amount of 5-minute epoxy to the top of the guide tubes, gluing the bundles in place. While the epoxy is curing, make sure that the wire bundles are still correctly aligned at the bottom.
9.7. To fix the moveable HPC wire arrays, first move the shuttle to the highest required position (in the experiments described in this article, at least 16 full turns/4mm above the lowest position). Then push the wire bundles into the u-shaped opening of the shuttle and glue them in place with a small amount of 5-minute epoxy. Make sure that the epoxy does not run down the bundle into the polyimide tube. When the epoxy is cured, apply a second layer of epoxy at the same spot to reinforce the connection and make it less likely for the connection to break when the shuttle is moved.
9.8. Carefully insert the open end of the wire-pin assembly of a GND wire through one of the through-holes marked GND. Fixate it using a gold pin. Note: When using a head stage in which GND and reference (REF) channels are shorted, a REF channel can also be used if more convenient.
9.9. Remove the drive from the holder, take care not to bend any of the wire assemblies. Re-attach the drive’s front part in the same holder and insert 4 EEG wire-pin assemblies into the through-holes for the EEG channels (marked 2,4, 29, 31) and fix them with a gold pin each.
9.10. For all GND and EEG wires, use a multimeter on the continuity setting to verify the continuous connection between the gold pin on the EIB and the pin of the connected wire-pin assembly.
9.11. Store the drive. This can be done for example by attaching the cap to the drive body and storing it upside down. NOTE: Prior to surgical implantation, sterilize the bottom of the drive using ethanol (see Table of Materials). All bone screws and GND/EEG wire assemblies should be sterilized in ethanol. Surgical instruments should be sterilized via an autoclave.

### 10. Drive implant surgery

NOTE: Section 10 briefly outlines the surgical procedures to implant the TD drive. A more extensive implantation protocol, including description of tools, as well as doses and concentrations of drugs can be found in the supplementary materials.

**Figure 2:**
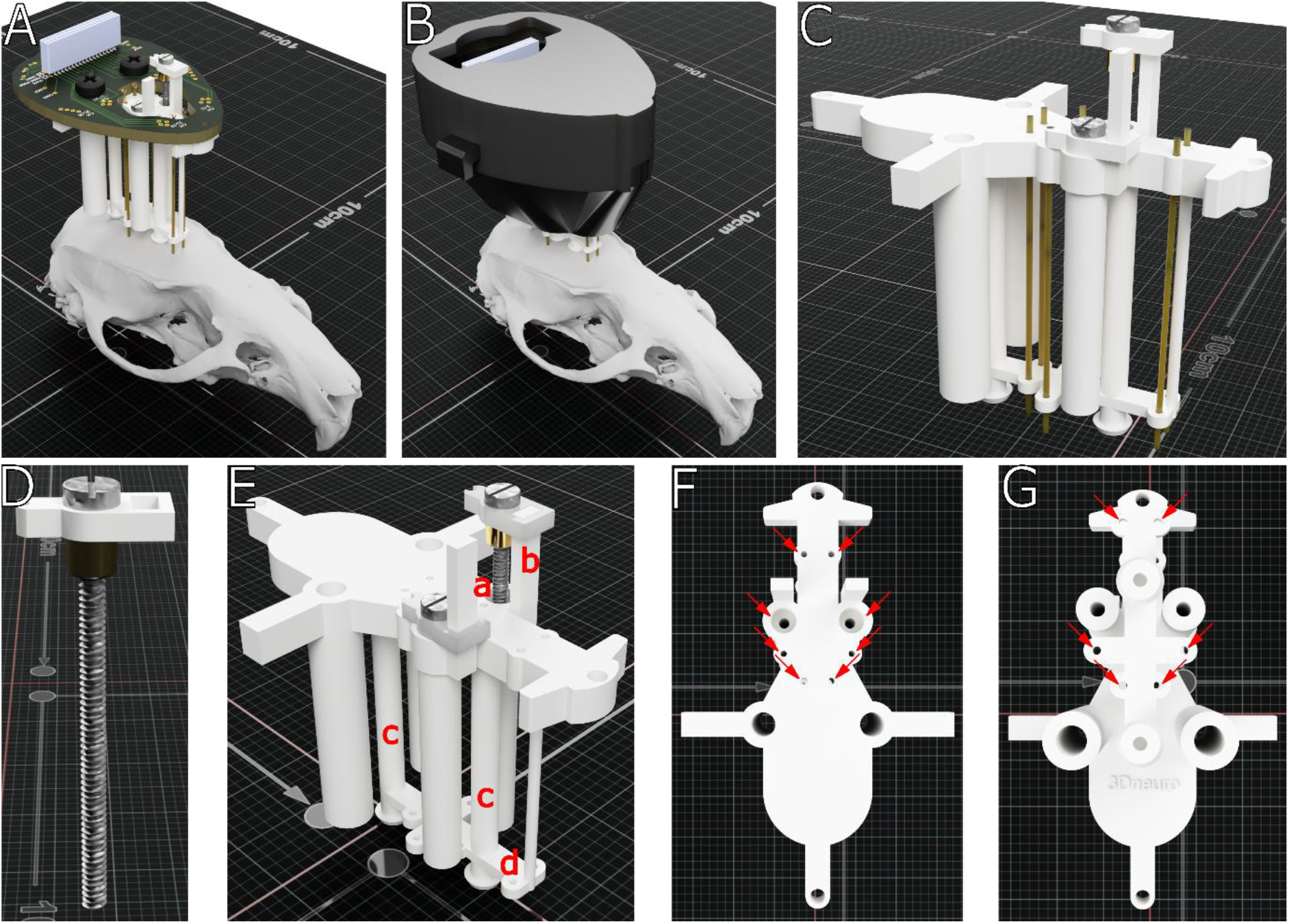
Rendering of the TD drive. **A & B**. TD drive with (A) and without protective cap on a rat skull model. **C**. Polyimide guide tubes correctly inserted into each of the six recording sites. **D**. An isolated, completed shuttle assembly, featuring the guide screw, 3D printed shuttle and the soldered brass insert. **E**. TD drive body with two shuttles inserted. Marked in red: **a** countersink holes for shuttle, **b** shuttle guide, **c** center pedestals of the drive body, **d** guide stencil **F & G**. Important locations on the top (F) and bottom (G) of the drive body that might require post-processing after 3D printing are indicated by a red arrow each.

10.1. Sterilize surgical tools and implant, clean and sanitize the surgical area.
10.2. Provide necessary pre-emptive analgesia, antibioticsm and gas anesthesia (isoflurane) in accordance with your local guidelines.
10.3. Place rat in the stereotactic apparatus. Shave the top of the head and disinfect the skin with povidone-iodine. Subcutaneously apply local anesthetic (lidocaine) and make an incision on the skull on top of the midline.
10.4. Expose the skull by pulling the skin to the side. Remove connective tissue on top of the skull, dry and clean the skull surface. Gently detach the muscles on the side of the skull to allow placement of anchoring screws.
10.5. Measure bregma and lambda coordinates. For precise targeting, ensure that the skull surface is parallel to the anterioposterior-mediolateral plane of the stereotax by measuring the difference in dorsoventral position of bregma and lambda. If the coordinates differ, adjust the position of the rat in stereotax by raising of lowering the mouth piece.
10.6. Mark craniotomies around the target locations (prelimbic cortex (AP +3.5 mm and ML + -1 mm), retrosplenial (AP+5.8 mm and ML +-1mm and hippocampus (AP -3.8 mm and ML + - 2.5 mm))
10.7. Drill holes for GND/EEG screws and anchoring screws. Insert the screws and cover them with liquid dental acryllic. Drill the craniotomies and carefully remove the dura mater. Prevent craniotomies from drying out by applying sterile saline.
10.8. Carefully position the TD drive on top of the craniotomies, making sure that the guide tubes are flush with the skull. Protect the guide tubes with petroleum jelly and attach the TD drive to the skull with dental acrylic.
10.9. Lower the wire arrays targeting HPC slowly from their intial location (∼1.5mm DV from brain surface) towards the pyramidal layer of hippocampal CA1. The pyramidal layer was reached progressively in the subsequent days during signal checks in the rats’ recovery period.
10.10. Place the protective cap around the drive.
10.11. Turn off gas anethesia and remove rat from the stereotactc frame. Place the rat in a clean cage inside a heated chamber and provide wet food and water for recovery. Monitor the rat until is active again, moving in the cage, eating and drinking.
10.12. Provide appropriate post-surgical care.

### 11. EIB recovery

11.1. After the end of the experiment, recover the drive and remove the protective cover.
11.2. Remove the gold pins and connected electrode wires carefully. Unscrew the EIB from the drive body. By gently pushing soft tweezers between the EIB and the drive body or carefully lifting the EIB by hand, release the remaining cyanoacrylate bond that holds the EIB to the body.
11.3. Clean the EIB and gold pins for reuse on subsequent TD drive implants. Before re-using an EIB, check the gold pin vias and high-density connector for wear. Only re-use the EIB if the vias are intact enough to allow good connection between gold pins, electrode wires, and EIB and if the high-density connector’s connection to the headstage is still sufficiently stable.

## REPRESENTATIVE RESULTS

Using the instructions provided in the protocol, the TD drive could be built by multiple experimenters (A.S., J.v.d.M., P.v.H.) easily. After drive-development (n=4) a full pilot was ran with eight animals. An additional batch of eight animals has been implanted and underwent experimental data collection. As data analysis has not been completed on these animals, they have been included in the survival analysis, but not in other analyses (e.g., targeting or histology).

Implant surgery was performed 2 weeks after arrival (see Figure 3A for the target locations used in the pilot). The implant was performed with usual surgical procedures (see above) and lasted ∼3h. An experienced surgeon (L.G.) performed initial implants and could teach both experienced as well as novice experimenters with 2-3 surgeries to independence.

**Figure 3:**
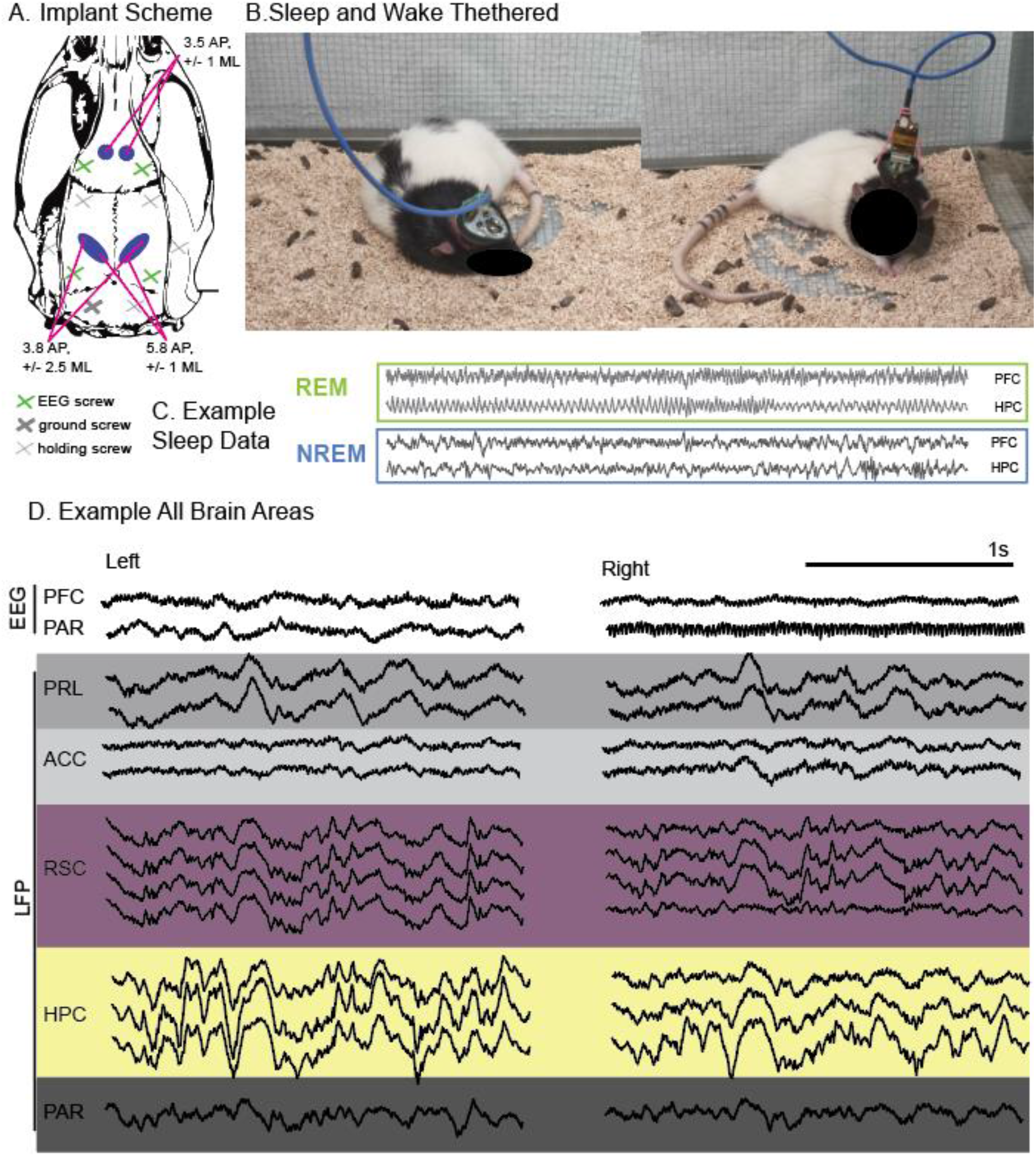
**A**. Schematic overview showing the target locations for craniotomies (blue circles) and skull screws (green: EEG, blue: GND, gray: anchoring screws, note that two anchoring screws are on the side of the skull). **B**. Photo of implanted animals with a tethered headstage during sleep and wakefulness. **C**. Example sleep data from tethered animal PFC (Prelimbic) and HPC (Ca1), divided in REM sleep with theta and NonREM sleep with delta, spindles and ripples. Y axis microvol, x axis seconds. This data can be used for example for sleep scoring or oscillation event detection and analysis **D**. Example broadband activity recorded wirelessly in an awake animal (noisy channels on the left were not connected).

All animals recovered well and tolerated the implant (Figure 3B). Frontal and retrosplenial electrodes were fixed, but the hippocampal bundles were movable. Hippocampal bundles were implanted at a dorsoventral depth of 2mm and adjusted to maximize HPC coverage during two weeks of surgery recovery, where the signal was checked live during sleep habituation periods. In 7 out of 8 animals, all target sites were reached on at least one hemisphere (Table 2 for hit rates, see Figure 4C for representative histology). Wake and sleep recordings were performed successfully tethered in a recording box as well as wireless recordings in a larger maze (example data Figure 3C and D). Animals kept the implants for 2 months, when individual animals would start losing them, however majority of animals kept the implants until experimental end day 85-100 after implant (Figure 4A). During this time the LFP remained stable as can be seen in an example analysis where we detected delta oscillations (Figure 4B). There was normal variability across time but no systematic drift of the signal in any of the recorded brain areas (including pyramidal layer of CA1). It is recommended to end experiments within 10-15 weeks after surgery. All EIBs could be recovered.

**Table 1:**
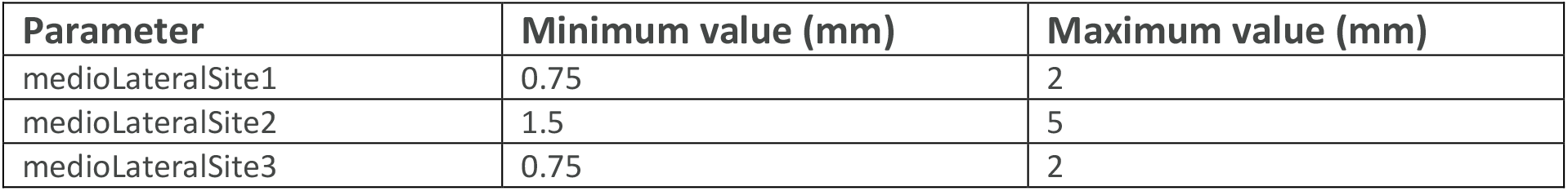
Overview of the manually imposed limits on the parameters controlling the mediolateral coordinates of the recording sites.

| Parameter | Minimum value (mm) | Maximum value (mm) |
| --- | --- | --- |
| medioLateralSite1 | 0.75 | 2 |
| medioLateralSite2 | 1.5 | 5 |
| medioLateralSite3 | 0.75 | 2 |

**Table 2:** Hit rate for pilot of 8 animals. In 4 out of 8 animals, all electrodes were placed correctly. However, in 7 out of 8 animals, all brain areas were correctly targeted in at least one hemisphere (with the exception of 1 rat missing CA1 pyramidal layer).

| Hemisphere | PFC | RSC | CA1 pyr. |
| --- | --- | --- | --- |
| Right | 8 of 8 | 5 of 8 | 6 of 8 |
| Left | 8 of 8 | 7 of 8 | 7 of 8 |

**Figure 4:**
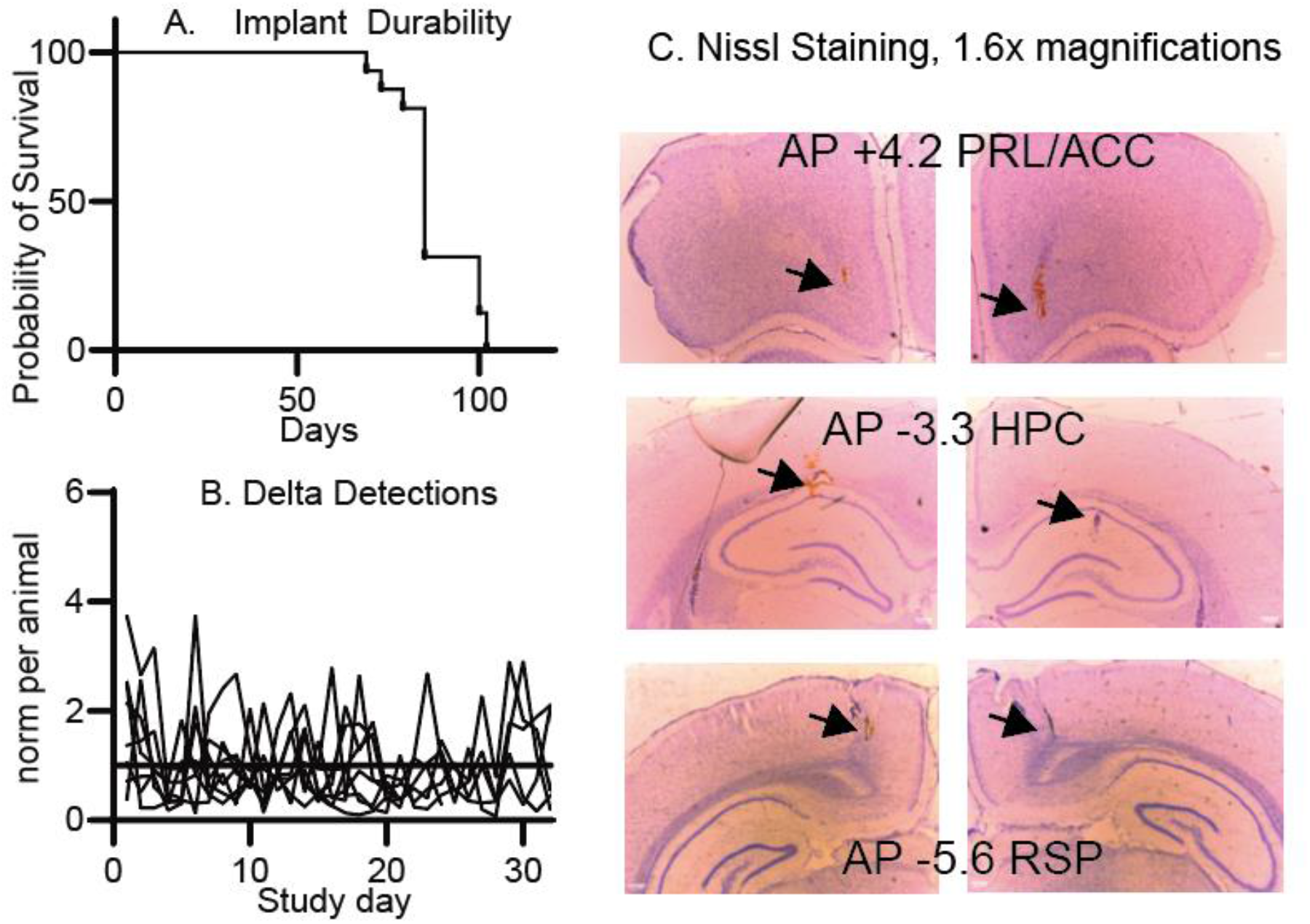
A. Survival plot for implant for two rounds of longer running experiments. Of note, on day 85 n=8 finished experiment and were planned perfused. B. Data example for stability. Shown are the count of delta detections in the hippocampal channel over recording days (∼3 days per week). Each animal showed a normal variation depending on the amount of sleep but there was no general drift over time in the signal and thus detections.

We have applied this implant mainly to measure 1, sleep stages and 2, sleep oscillations in response to learning and other interventions. For example, how oral CBD intake influences oscillation occurrence and coherence across brain areas (see ^20^)

## DISCUSSION

Presented in this article, is an adaptable implant for bilateral, symmetric multi-area wire electrode recordings for freely-moving rats.

The ability to easily adjust the implant by changing predefined parameters was one of the motivations for the creation of the TD drive. While aiming to maximize the flexibility for changing parameters, inherent constraints in the relations between them necessarily imposed limits to this adaptability. No limits are set by default for the antero-posterior parameters, with these coordinates instead being governed by logical inter-site interactions and the overall size of the drive body. The parameters controlling the mediolateral coordinates of all recording sites are subject to manually imposed limits (see Table 1).

Underlying the many parametric options are numerous interactions between the design features. These interactions can become disjointed under certain conditions. In an ideal situation, all possible combinations of parameter values are valid. However, with the more complex design of the TD Drive, it was opted to limit the mediolateral coordinates to within the tested range. Chosing coordinates outside the tested limits is, in principle, possible. However, making such changes is not recommended as the integrity of the design can become compromised and restoring it while maintaining the out-of-limit coordinates might require CAD modelling experience. This is an inherent result of the trade-off between ease of drive production and flexibility – larger degrees of freedom do make the parametric definition of the drive more complex and can result in overly fine-grained, undesired outcomes (for example, the need to produce different EIBs for small changes in the drive design). The parameter choices made in this manifestation of the TD drive are guided by preferences of the current experimenters. It was opted, for example, to increase the height of the TD drive’s pedestals, improving ease of implantation during surgery at the cost of a slightly higher final implant. However, the implant is still much smaller than drives with individually moveable tetrodes and was well-accepted by the animals. The current design of the TD drive has been created in Autodesk Fusion 360. While it is one of the most advanced programs for parametric 3D computer-aided design and at time of publication does provide a free license for academic use, the commercial and cloud-based nature of the program does pose a risk for the free availability of the design. Therefore, porting the design to a true open source parametric CAD software^21^ such as FreeCAD might be necessessary for a future iteration.

The surgery for the TD drive can be performed in 2-3 h. The target locations of the wire are marked stereotactically (PFC +3.5AP next to midline, HPC –3.8AP -/+ 2.5 ML, RSC –5.8 next to midline) and the screw electrodes, GND, and additional skull screws for stability are placed relative to those locations. While the TD drive stencil does provide stereotactic locations, imprecisions in placement of the polyimide tube and the use of less stiff tubing material can introduce small variations in the position of the electrodes. Therefore, it is recommended to drill small craniotomies (instead of guide tube-sized 0.5mm burr holes) to account for this variability. In our surgery, a single, larger craniotomy for the RSC and HPC targets is drilled. For PFC and RSC, it was chosen to have wire bundles implanted at a fixed depth. The PFC bundle had wires targeted at two different depths to record from the prelimbic as well as the anterior cingulate cortex. The HPC bundles were movable and were built with 3 wires at varying heights to facilitate reaching the Ca1 pyramidal layer as well as the stratum radiatum. The last, shorter wire allowed recording from the PPC. We achieved the best results for targeting the hippocampal Ca1 when the longest electrode wire was moved to the target depth (2mm ventrally from the brain surface) during surgery time, with only small adjustments in the two weeks after surgery during live-signal checks to accommodate for individual variances and brain swelling after surgery.

A problem with large implants, such as tetrode hyperdrives, in rats is the chance of the implant stability degrading and animals losing the implant. For the TD drive individual failures were observed due to degradation of implant stability after 2 months (3 out of 16 animals). Therefore, the TD drive is recommended for experiments with an intended maximum duration of 10-15 weeks. We show that for this time period the signal is stable – even the hippocampal pyramidal layer recordings that are precise – and there is no systematic drift or significant wobble affecting the LFP recordings.

With a total assembly time of around 3 hours and surgery time of around 2-3 hours, the TD drive offers a compromise between tetrode hyperdrives and simpler, less adjustable multi-wire implants^22^. With our chosen targets, recordings from 10 brain areas with 6 bundles were achieved. Compared to other implants for non-moveable wire bundles, the symmetric placement of recording sites yields another advantage: if lateralization is not relevant, simultaneously implanting wires in both hemispheres increases the chance of hitting the correct target and therefore also the data yield per animal. In the pilot, 4 out of 8 animals had all 5 target sites (PFC with Prl and ACC, RSC, PPC and Ca1 pyramidal layer of HPC) targeted correctly bilaterally, but 7 out of 8 had at least each brain area on one side recorded. Thus, this drive is advisable for those that want a quick and easy-build solution to record LFP that can be applied by less experienced researchers especially when higher animal numbers are needed such as in sleep studies. With many high-end tetrode hyperdrives, that would allow for the recording of individual neuronal activity, even very skilled and experienced researchers can only build and implant 2-6 implants per year of which many will not successfully reach any brain area of interest. It takes many years of training to achieve higher success rates and even then, the number of animals that can be recorded efficiently remains low.

In sum, the TD drive presents an easy and fast to build wire-drive with 6 bundles that can be easily adapted to contain different recording sites and other implants such as cannulas and fibers.

## ACKNOWLEDGMENTS

The authors would like to thank Angela Gomez Fonseca for the inspiration to develop the drive and all students that ran pilot experiments with the animals, Milan Bogers, Floor van Ravenswoud, Eva Severijnen. This work was supported by the Dutch Research Council (NWO; Crossover Program 17619 “INTENSE”).

