## Supplemental figure 1 for "The TD drive - A parametric, open-source implant for multi-area electrophysiological recordings in behaving and sleeping rats"

**S1**

**Channel mapping for use with Intan RHD 32 channel headstage, Intan chip (Omnetics writing) on headstage facing to front of TD Drive. Numbers in white correspond to the mapped channels in the data**

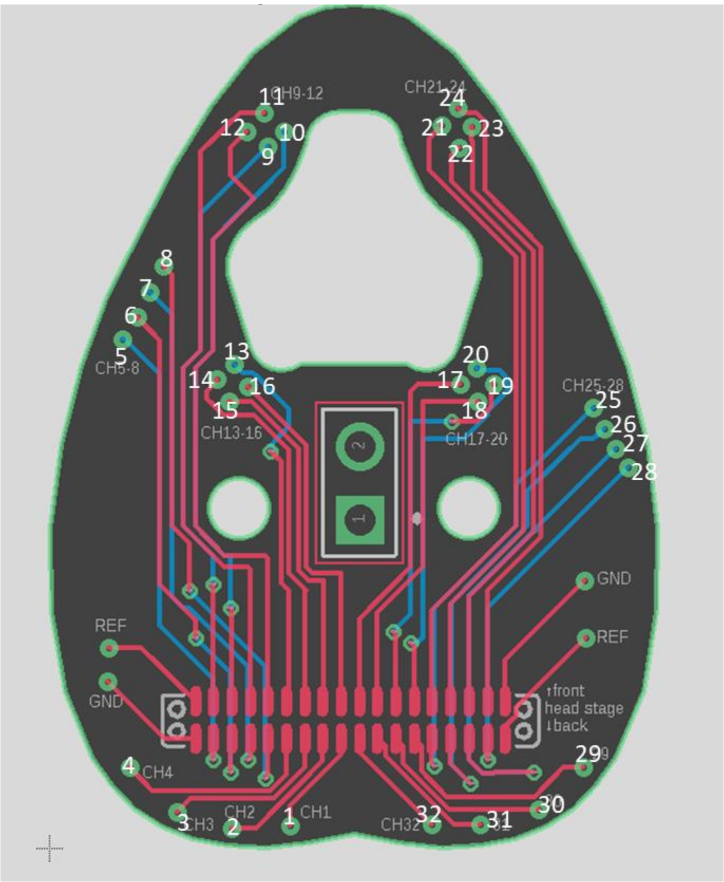


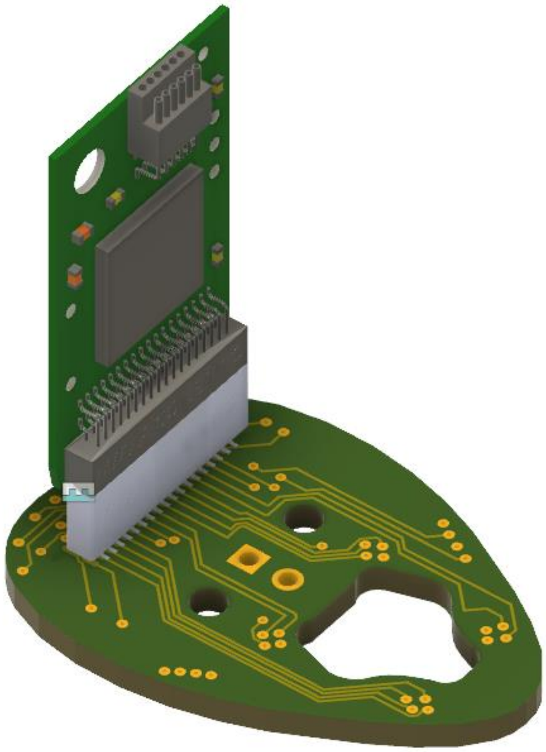

**Top view of the EIB with schematic of the different wire bundle configurations used.**

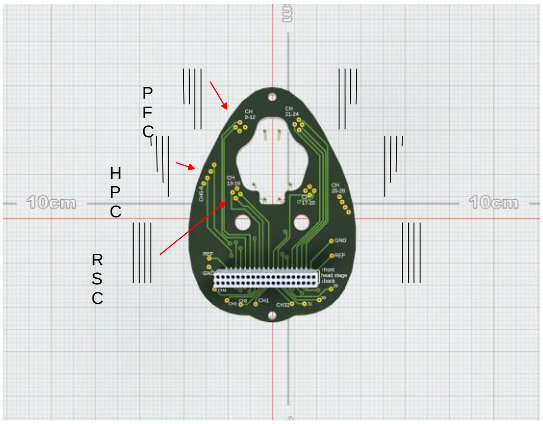


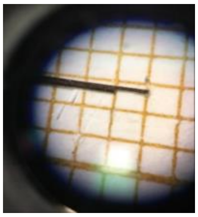

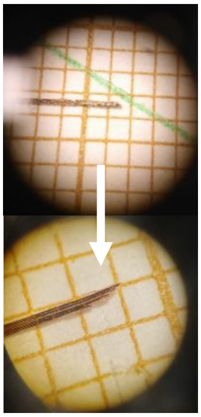
