## Supplemental figure 2 for "The TD drive - A parametric, open-source implant for multi-area electrophysiological recordings in behaving and sleeping rats"

## S2
### Additional pictures showing intermediate drive building progress

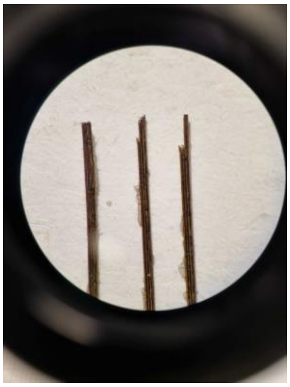

**(a) Additional view of the three wire bundle ends. Left: RSC, center: HPC, right: PFC**

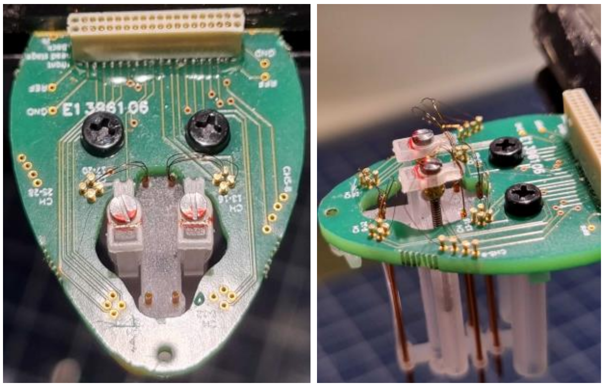

**(b) Left: Top view of a drive with RSC wires inserted. Right: side view of a drive with PFC, HPC, and RSC wires connected to the EIB**

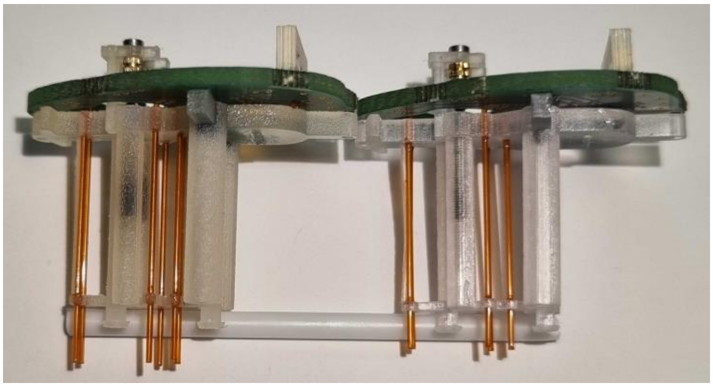

**(c) Left: TD drive with guide tubes of varying length at the bottom. Right: TD drive with polyimide tubes cut to be flush with the brain at implantation.**

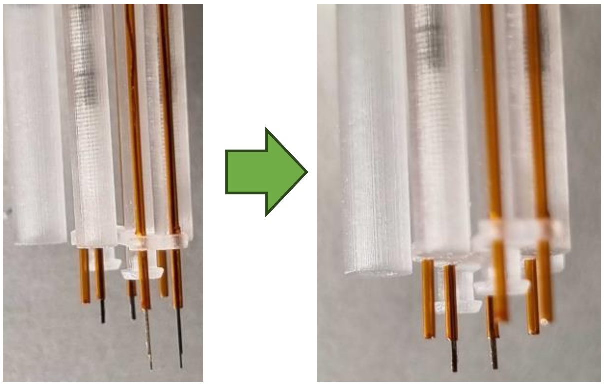

**(d) Adjust the length of ire bundles to correctly target the recording locations (see Step 9.5 in the main protocol).**
